# Defective excitatory/inhibitory synaptic balance and increased neuron apoptosis in a zebrafish model of Dravet syndrome

**DOI:** 10.1101/781393

**Authors:** Alexandre Brenet, Rahma Hassan-Abdi, Julie Somkhit, Constantin Yanicostas, Nadia Soussi-Yanicostas

## Abstract

Dravet syndrome is a type of severe childhood epilepsy that responds poorly to current anti-epileptic drugs. In recent years, zebrafish disease models with Scn1Lab sodium channel deficiency have been generated to seek novel anti-epileptic drug candidates, some of which are currently undergoing clinical trials. However, the spectrum of neuronal deficits observed following Scn1Lab depletion in zebrafish larvae has not yet been fully explored. To fill this gap and gain a better understanding of the mechanisms underlying neuron hyperexcitation in Scn1Lab-depleted larvae, we analyzed neuron activity *in vivo* using combined local field potential recording and transient calcium uptake imaging, studied the distribution of excitatory and inhibitory synapses and neurons as well as investigated neuron apoptosis. We found that Scn1Lab-depleted larvae displayed recurrent epileptiform seizure events, associating massive synchronous calcium uptakes and ictal-like local field potential bursts. Scn1Lab-depletion also caused a dramatic shift in the neuronal and synaptic balance toward excitation and increased neuronal death. Our results thus provide *in vivo* evidence suggesting that Scn1Lab loss of function causes neuron hyperexcitation as the result of disturbed synaptic balance and increased neuronal apoptosis.

## 1. Introduction

Dravet syndrome (DS) is a type of severe drug-resistant childhood epilepsy. It is associated with several comorbidities, including intellectual disability, irreversible psychomotor deficits, behavioral disorders, and high mortality at around 10 years of age [1], [2]. DS is a genetic disease, caused in 80% of cases by de novo loss-of-function mutations in the *SCN1A* gene encoding the alpha-1 subunit of the main voltage-dependent sodium channel in inhibitory interneurons [3], whose synapses mainly release GABA, a neurotransmitter that inhibits excitation of post-synaptic neurons [4].

Among animal models that have been developed in recent years, the zebrafish has proved to be a versatile and powerful system for *in vivo* epilepsy research [5]–[7]. In particular, zebrafish larvae with loss of function of the *scn1Lab* gene, one of the two zebrafish orthologs of *SCN1A*, have already been used as animal models to find novel antiepileptic drug candidates, several of which are now undergoing clinical trials [8]–[10].

However, while Scn1Lab-depleted larvae have been established as a *bona fide* epilepsy model [7]–[15], the spectrum of neuronal deficits induced by the loss of function of this sodium channel has not yet been fully explored. To fill this gap, we first combined transient calcium uptake imaging and local field potential recording to analyze neuron hyperactivity at whole brain levels in real time. In Scn1Lab-depleted larvae, we observed recurrent events associating massive synchronous calcium uptakes and ictal-like local field potential bursts characteristic of epileptiform seizures. Importantly, we also observed that these larvae displayed a marked deficit in the number of inhibitory synapses, together with a dramatic over-accumulation of excitatory ones, heavily disrupting the synaptic balance, which was correlated to a shift in the neuronal population toward excitatory neurons. Last, we observed that loss of function of this sodium channel caused increased neuronal apoptosis. Taken together, our findings are evidence that Scn1Lab-depletion causes epileptiform seizures as a result of brain development defects that include impaired synaptic balance and increased neuronal death.

## 2. Materials and Methods

### 2.1 Fish maintenance and transgenic lines

Adult zebrafish were raised at 26.5 °C in standard aquaculture conditions with a 14 h light/10 h dark cycle. Embryos collected by natural spawning were raised on E3 medium complemented with 0.002% methylene blue and kept at 28.5 °C. 0.003% phenylthiourea (PTU; Sigma) was added at 1 day post-fertilization (dpf) to inhibit the pigmentation. The mutant *scn1Lab* (didy^S552^) was a gift from Dr. Herwing Baier (Max Planck Institute of Neurobiology, Germany), the HuC:GCaMP5G transgenic line was a gift from Dr. George Debrégeas (Laboratoire Jean Perrin, Paris) and the Gad1b:GFP; Vglut2a:DsRed double transgenic line was a gift from Dr Germán Sumbre (IBENS, Paris).

All the animal experiments described in the present study were conducted at the French National Institute of Health and Medical Research (INSERM) UMR 1141 in Paris in accordance with European Union guidelines for the handling of laboratory animals (http://ec.europa.eu/environment/chemicals/lab_animals/home_en.htm), and were approved by the Direction Départementale de la Protection des Populations de Paris and the French Animal Ethics Committee under reference No. 2012-15/676-0069.

### 2.2 Morpholino

Antisense morpholino oligonucleotide (5’-CTGAGCAGCCATATTGACATCCTGC-3’), obtained from Gene Tools, was used to block the *scn1Lab* zebrafish mRNA transcription. One-to two-cell embryos were injected with 1 pmol MO scn1Lab^AUG^, 0.53 ng rhodamine B dextran and 0.1 mM KCl.

### 2.3 Locomotor activity

Larvae locomotor activity (*i.e.* movement) was evaluated using the Zebrabox, an infrared automated recording and tracking device supported by ZebraLab software. Each 96-well plate containing 4 dpf control, morphant or mutant larvae in 200 µL E3 medium was placed in the Zebrabox recording chamber. In all locomotion recording protocols, animal color was set to dark and detection threshold to 15. After 45 min habituation in darkness, larvae were simultaneously tracked for 25 min. Larvae movement in each well was computed as the sum of all pixels whose intensity changed during the recording, and plotted as “acting units”.

### 2.4 Calcium imaging

4 dpf zebrafish larvae were paralyzed using 300 µM pancuronium bromide (PB, Sigma) and immobilized upside down in a recording chamber in 1.2% low-melting agarose covered with E3 medium containing 0.003% PTU and 300 µM PB. The chamber was then placed under a Leica SP8 laser scanning confocal microscope equipped with a 20x/multi-immersion 0.75 objective. Calcium uptake events were detected by recording the fluorescence of a 512 × 512 pixel image of a single focal plane at 2 Hz for 1 h. Fluorescence intensity of the optic tectum was measured using ImageJ software. Fluorescence variations (*ΔF/F_0_*) were calculated by subtracting the mean fluorescence intensity of all frames and normalizing the mean fluorescence intensity of all frames. The fluorescence drift over time was corrected by subtracting the mean value of the lowest values (all values under the median) within a 20 s sliding window around the point. All fluorescence increases greater than 0.04 *ΔF/F_0_* were considered as calcium events. Since the detection system may detect false events, all of them were manually checked.

### 2.5 Local field potential recording

4 dpf zebrafish larvae were paralyzed using 300 µM PB and immobilized, central side down, in 2% low-melting agarose covered with E3 medium containing 300 µM PB. A glass electrode (5 - 6 MΩ) filled with artificial cerebrospinal fluid composed of 10 mM HEPES, 134 mM NaCl, 2.9 mM KCl, 2.1 mM CaCl_2_, 1.2 mM MgCl_2_, 10 mM glucose; pH 7.8, was placed in the left neuropil of the optic tectum of the larva. The recordings were performed for 1 h in a current clamp mode at 10 µs sampling interval and with a 0.1 Hz high-pass filter, 1 kHz low-pass filter and digital gain at 10 (MultiClamp 700B amplifier, Digidata 1400 digitizer, both Molecular Devices). Results were analyzed with the Clampfit software (Molecular Devices). Events were defined as every downward membrane potential variation under −0.3 mV amplitude and lasting more than 100 ms.

### 2.6 Synapse immunostaining

4 dpf control and morphant larvae were fixed with 4% formaldehyde for 1 h 30 min at room temperature and stored at 4 °C in PBS containing 0.02% sodium azide. Fixed larvae were transferred to 15% sucrose at 4 °C overnight and then embedded in 7.5% gelatin/15% sucrose, flash frozen in isopentane at −45°C and stored at −80 °C. When needed, frozen embedded larvae were cut into 20 µm thick sections using a cryostat, mounted on Superfrost slides and stored at −20°C until immunostaining. For gephyrin, larvae were not fixed but transferred in 15% sucrose for 1 h at room temperature, anesthetized with 0.01% tricaine and embedded in 7.5% gelatin/15% sucrose. 20 µm thick sections were post-fixed for 10 min with 4% formaldehyde and treated with 0.25% trypsin diluted in PBS for 2 min. Immunostaining was performed as follows. Sections were washed three times with PBS, blocked and permeabilised with 0.2% gelatin/0.25% Triton diluted in PBS. Rabbit antibody anti-PSD-95 (Abcam, Ab18258, 1:200), an excitatory post-synaptic protein, or rabbit antibody anti-gephyrin (Abcam, Ab185993, 1:100), an inhibitory post-synaptic protein, were added onto the sections, which were incubated overnight at room temperature. Sections were washed several times and incubated with anti-rabbit IgG Alexa 488 conjugated antibody (Molecular Probes, A-21206, 1:500), in the dark at room temperature. After 1 h incubation, sections were washed and stained in the dark for 10 min with 0.3% DAPI. After a last wash, sections were mounted in Fluoromount medium and conserved at 4 °C in the dark. Sections were imaged using a Leica SP8 laser scanning confocal microscope equipped with a 40x/oil 1.3 objective. Optic tectum and telencephalon regions were captured at full resolution, 0.063 × 0.063 × 0.4 µm voxel size. Images were deconvoluted using AutoQuant (Media Cybernetics) software and processed using ImageJ software. PSD-95 and gephyrin puncta were quantified using a semi-automated macro (Zsolt Csaba; Inserm UMR1141), which applied a threshold to segment the image and select only the puncta ranging between 0.018 µm^2^ and 3.14 µm^2^. The total number of PSD-95 or gephyrin puncta was finally divided by the surface are of interest to obtain the density of PSD-95 and gephyrin signal in the optic tectum. All manual cell counts were performed blind.

### 2.7 Identification and counting of inhibitory and excitatory neurons

3, 4 and 5 dpf larvae were anesthetized using 0.01% tricaine and embedded in 1.2% low-melting agarose in a recording chamber covered with E3 medium containing 0.003% PTU and 0.01% tricaine. The chamber was then placed under a Zeiss LSM 880 laser scanning confocal microscope equipped with a WPApo 20x/1.0 objective. 52 slices spanning approximately 100 µm were captured and processed using AutoQuant 3.1X (Media Cybernetics). Gad1b and Vglut2a positive neurons in the optic tectum were counted using Imaris MeasurementPro8.4.2 (Bitplane). Briefly, neurons were approximated as 7 µm diameter spheres and sorted by their fluorescence intensity.

### 2.8 Neuronal death

Neuronal death was detected in living zebrafish brains using acridine orange (AO), which binds to free nucleic acid. 4 dpf control and *scn1Lab* larvae were incubated for 20 min with 10 µM AO (VectaCell) in E3 medium. Larvae were washed several times, paralyzed with 300 µM PB and embedded in 1.1% low-melting agarose in the center of a 35 mm glass-bottom dish, covered with E3 medium containing 0.003% PTU and 300 µM PB. 100 µm brain stacks were acquired using a Leica SP8 laser confocal scanning microscope equipped with a 20x/multi-immersion 0.75 objective. In addition, 4 dpf control and morphant brains were dissected, fixed with 4% formaldehyde and stained with activated caspase-3 antibody. The brains were first washed with 0.1% triton in PBS, then permeabilised and blocked with 5% goat serum/1% triton in PBS. Rabbit polyclonal antibody to activated caspase-3 (Abcam, Ab44976, 1:500) was added, which was incubated overnight at room temperature. Brains were washed several times and incubated with an anti-rabbit IgG Alexa 488 conjugated antibody (Molecular Probes, A-21206 1:500), for 1 h at room temperature in the dark. Brains were finally washed and embedded in 1.1% agarose in the center of a 35 mm glass-bottom dish and imaged using a Leica SP8 laser scanning confocal microscope equipped with a 20x/multi-immersion 0.75 objective. All images were processed using AutoQuant 3.1X (Media Cybernetics) and ImageJ software.

### 2.9 Colocalization of activated caspase-3 labeling with inhibitory or excitaroty neurons

4 dpf Tg[Gad1:GFP; Vglut2a:DsRed] larval brains were stained using activated caspase-3 antibody followed by anti-rabbit IgG Cy5 conjugated antibody (Molecular Probes, A-10523,1:300) and imaged as previously described. Images were processed using AutoQuant 3.1X (Media Cybernetics). Colocalization of activated caspase-3 signals with Gad1b or Vglut2a positive neurons was assessed manually using ImageJ software.

### 2.10 Z-VAD treatment

3 dpf control and *scn1Lab* larvae were incubated for 24 h with either vehicle or 300 µM Z-VAD-fmk (Sigma), in E3 medium containing 1% dimethyl sulfoxide (DMSO). Dead cells were then labeled using AO and imaged as previously described.

### 2.11 Statistics

Data were plotted and analyzed using GraphPad Prism 5. Data were checked for normality using the d’Agostino and Pearson omnibus normality test with alpha set at 0.05. Data that did not follow a normal distribution were compared with a Mann-Whitney test. Data that followed a normal distribution were analyzed with Student’s two-tailed unpaired *t-test* with or without the Welch correction depending on the variance difference of each sample. Graphs show the mean and the SEM.

## 3. Results

### 3.1 Calcium imaging combined with LFP recordings as tools to analyze neuronal activity in vivo in Scn1Lab-depleted zebrafish larvae

As previously described, zebrafish embryos injected with 1 pmol *scn1Lab*^AUG^ morpholino-oligonucleotide, hereafter referred to as *scn1Lab* morphants, and *scn1Lab^552/s552^* mutants displayed several highly penetrant phenotypes, including black appearance resulting from defective pigment aggregation, failure to inflate their swim bladder, and, from 4 dpf onward, recurrent spontaneous epileptiform seizures, reflected by a markedly increased swimming activity of the larvae [8], [10], (Figure S1). Importantly, since phenotypes observed in *scn1Lab* morphants (*N* = 904), and particularly seizure number and intensity, were the same as those seen in *scn1Lab^s552/s552^* mutants (*N* = 192), we decided to use the former to further characterize the neuronal phenotypes associated with loss of Scn1Lab function.

First, to confirm that the increased swimming activity observed in 4 dpf *scn1Lab* morphants did correspond to *bona fide* epileptic seizures caused by the synchronous hyperexcitation of large neuron populations, we generated zebrafish *scn1Lab* morphants that also carried the Tg[HuC:GCaMP5G] transgene encoding a calcium-dependent fluorescent protein expressed under the control of the pan-neuronal HuC promoter [16]. To record transient calcium uptakes associated with neuron hyperexcitation during epileptiform seizures, we imaged the optic tectum of pancuronium-paralyzed and agarose-immobilized 4 dpf larvae and simultaneously recorded forebrain local field potentials (LFPs) (Figure 1A). Results show that scn1Lab^AUG^ morphants displayed recurrent spontaneous transient calcium uptake events, events with *ΔF/F_0_* > 0.04, (Figure 1E, F; *p* < 0.001) that also precisely corresponded to multi-spike ictal-like discharges (correlation coefficient *R*^*2*^ = 0.772) (Figure 1G, H), which were never seen in either non-injected larvae or control morphants (Figure 1D) (Figure S2). Thus in good agreement with previous results [8], [10], 4 dpf Scn1Lab-depleted zebrafish larvae displayed epileptiform seizures, which could be detected *in vivo* by both calcium imaging techniques and LFP recordings.

**Figure 1.**
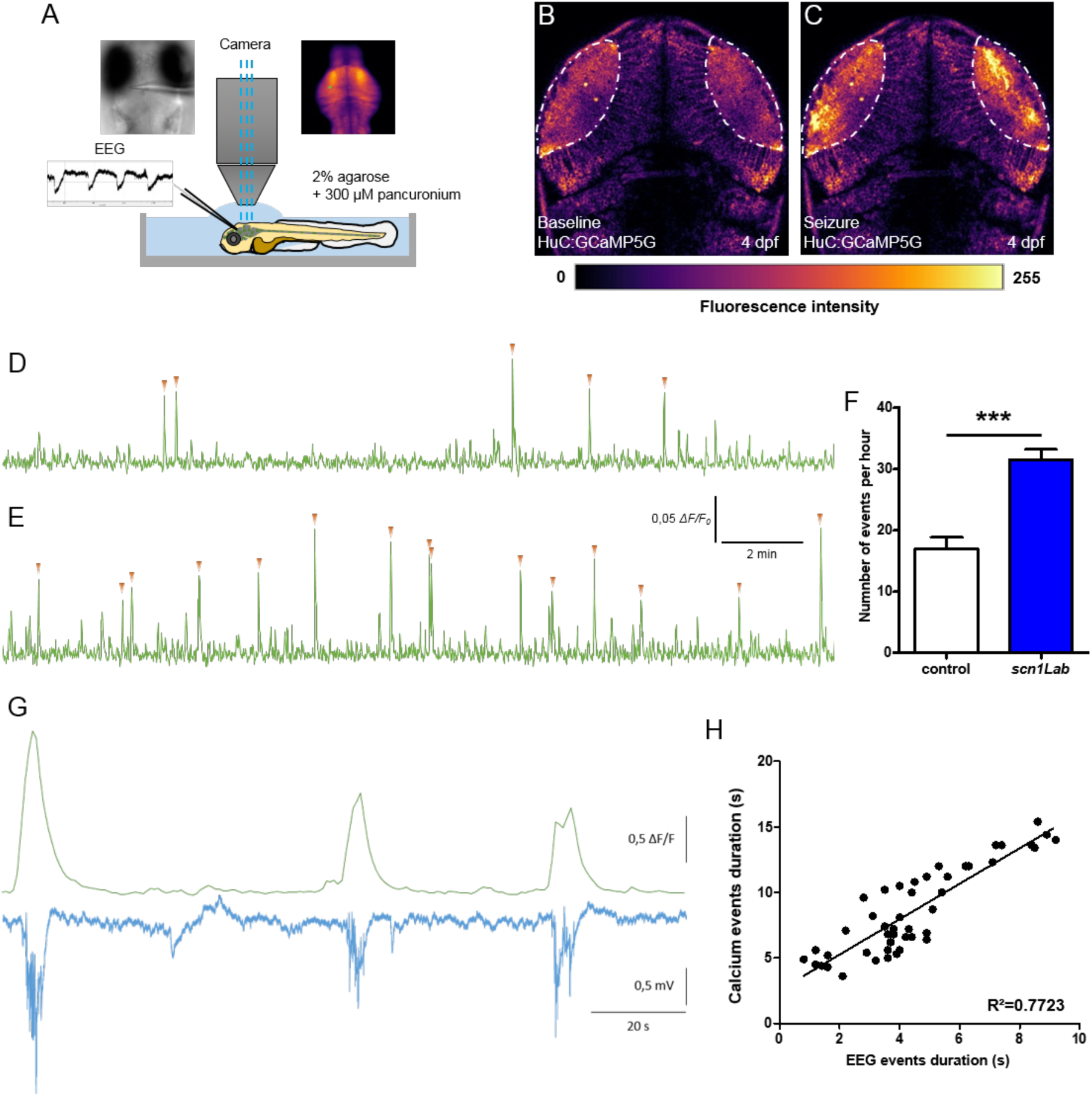
Correlation between LFP and calcium activity in *scn1Lab* model. (A) Schematic of the experimental setup to simultaneously record local field potential and calcium activity of a larval zebrafish brain. (B-C) Calcium images showing respectively the baseline activity (B) and the seizure activity (C). The fluorescence intensity is color-coded as shown in the color bar below the images. (D-E) Representative 20 min calcium recording of immobilized and paralyzed 4 dpf control (D) (*N* = 9) and *scn1Lab* (E) (*N* = 9) larvae. Each calcium event is shown with an orange arrowhead. (F) Number of events, calcium increase greater that 0.04 Δ*F/F*_0_ amplitude, during 1 h recording. (G) Representative LFP, recorded in the left neuropil of the optic tectum, and calcium traces, from the optic tectum, of a paralyzed and immobilized 4 dpf *scn1Lab* larva. (H) Correlation between LFP and calcium events duration (*n* = 46). *N* = number of larvae and *n* = number of events. Error bars on all graphs represent standard error of the mean (SEM). ***, *p* < 0.001. *p*-value was determined using Student’s unpaired *t*-test.

### 3.2 Inhibitory/excitatory synaptic balance is disrupted in Scn1Lab-depleted larvae

Harmonious functioning of neuron networks relies on a finely tuned balance between excitatory and inhibitory inputs [17]. Synapses, which are essential components of this balance, comprise both excitatory and inhibitory subunits that release glutamate and GABA, respectively [18]. This balance is impaired in neurological disorders such as epilepsy in which large populations of neurons display synchronous hyperexcitation [19], [20]. We recently showed that zebrafish *gabra1^-/-^*and *depdc5^-/-^* epileptic mutants displayed markedly fewer inhibitory synapses compared to age-matched control larvae, showing that these two mutants display brain development defects [21], [22]. To determine whether Scn1Lab deficiency also induced unbalanced excitatory/inhibitory post-synaptic outputs, we analyzed and compared, by immunocytochemistry, the distribution of the two post-synaptic populations on brain sections of 5 dpf control and Scn1Lab-depleted larvae, using anti-PSD-95 and anti-gephyrin antibodies, which label excitatory and inhibitory post-synaptic terminals, respectively [23]. Quantification of the two post-synaptic populations showed a marked increase in PSD-95 labeling (Figure 2B, C; *p* < 0.01) combined with a clear decrease of that of gephyrin (Figure 1E, F; *p* < 0.01), in scn1Lab^AUG^ morphants, compared to controls (Figure 2A, 2D). These data clearly show that depletion of Scn1Lab causes a marked disruption of the synaptic balance, which is shifted toward excitation (Figure 2G; *p* < 0.01).

**Figure 2.**
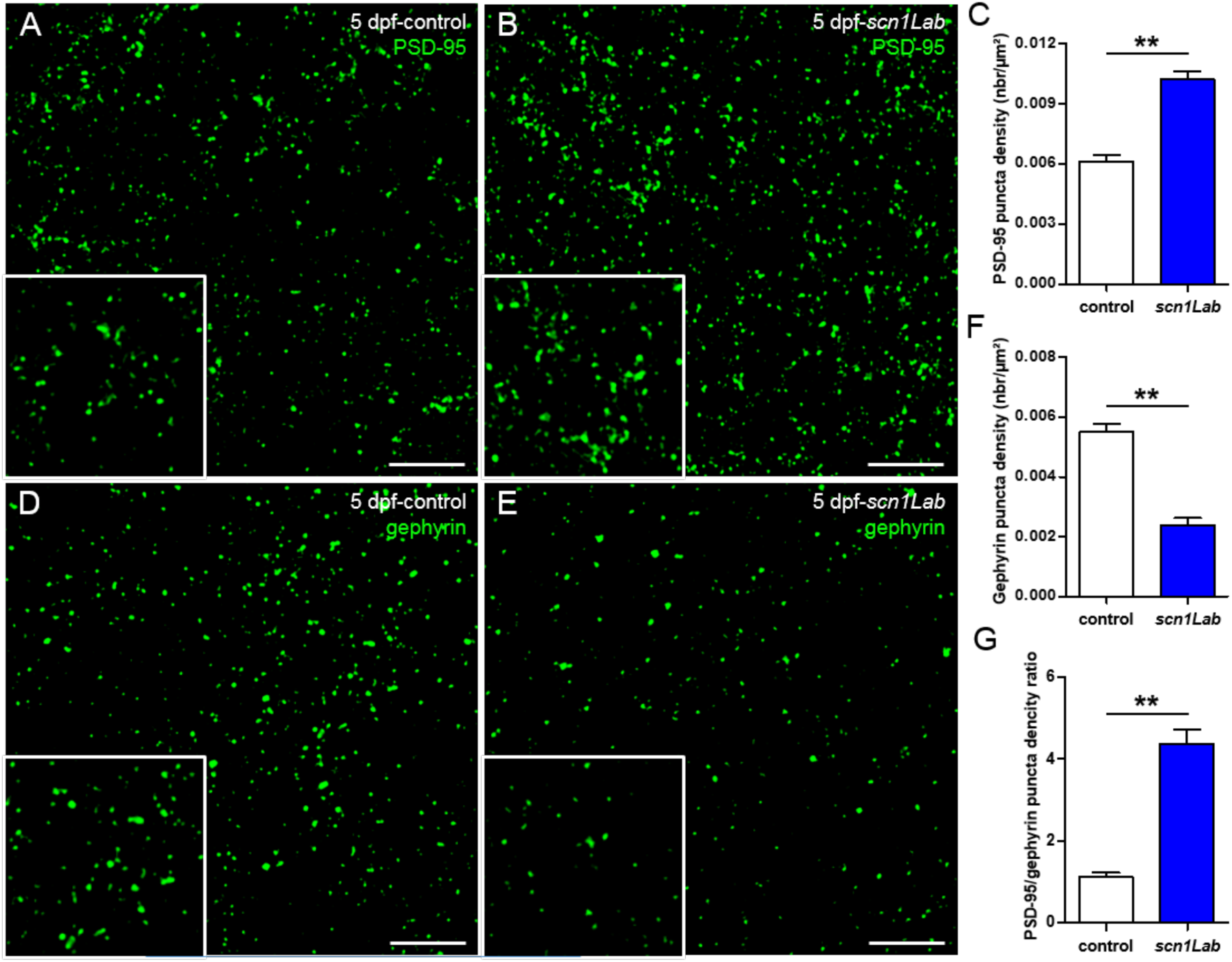
Defects of excitatory-inhibitory balance in the Scn1Lab*-*depleted larvae. (A-B) 20 µm coronal sections of 5 dpf control (A) (*N* = 5, *n* = 15) and *scn1Lab* (B) (*N* = 5, *n* = 15) larvae stained with PSD-95, an excitatory post-synaptic scaffolding protein. (D-E) 20 µm coronal sections of 5 dpf control (D) (*N* = 5, *n* = 20) and *scn1Lab* (E) (*N* = 5, *n* = 19) larvae stained with gephyrin, an inhibitory post-synaptic scaffolding protein. Scale bar 5 µm. Images were acquired with a SP8 Leica laser scanning confocal microscope equipped with a 40x/oil 1.3 objective. Scale bar 5 µm. (C, F) Quantification of PSD-95 (C) and gephyrin (F) puncta density. (G) PSD-95/gephyrin puncta density ratio. *N* = number of larvae and *n* = number of sections. Error bars on all graphs represent standard error mean (SEM). **, *p* < 0.01. *p*-values were determined using the Mann-Whitney test.

### 3.3 Distribution of excitatory and inhibitory neurons in scn1Lab model

We next assessed whether the defective excitatory/inhibitory synaptic balance seen in Scn1Lab-depleted larvae could be related to an abnormal ratio of inhibitory and excitatory neurons. To test this hypothesis, we used the double transgenic line Tg[Gad1b:GFP; Vglut2a:DsRed] that labels inhibitory neurons with GFP and excitatory neurons with DsRed [24], [25] as recipient line to generate scn1Lab^AUG^ morphants (Figure 3). First, live imaging combined with quantitative image analysis revealed that the total number of neurons is significantly decreased in scn1Lab^AUG^ morphants at 3 dpf (Figure 3D; *p* < 0.05), 4 dpf (Figure 3G; *p* < 0.001) and 5 dpf (Figure 3J; *p* < 0.01) when compared to controls. Next, we quantified the number of inhibitory (GFP^+^) and excitatory (DsRed^+^) neurons at 3, 4 and 5 dpf in scn1Lab^AUG^ morphants and controls. Our results show that at 3 dpf, the percentage of excitatory and inhibitory neurons was similar in Scn1Lab-depleted and control larvae (Figure 3K). By contrast, the ratio of excitatory to inhibitory neurons was markedly increased at 4 dpf (Figure 3L; *p* < 0.001) and 5 dpf (Figure 3M; *p* < 0.01). It is of note that the ratio of inhibitory to excitatory neurons did not change in control embryos between 3 and 5 dpf (Figure 3K, L, M).

**Figure 3.**
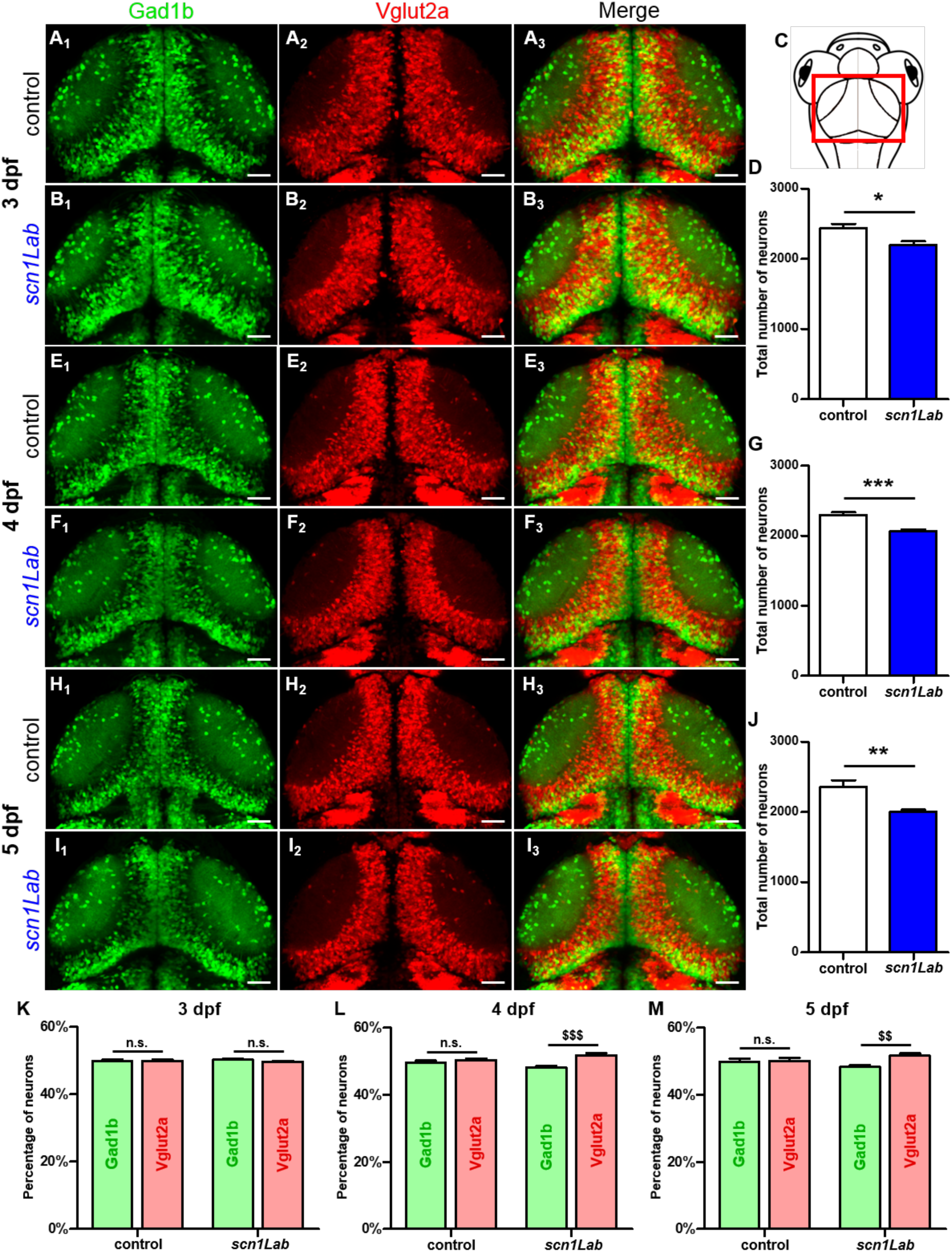
Evolution of the excitatory-inhibitory neuronal population in the *scn1Lab* model. (A_1-3_, B_1-3_) Dorsal view of 3 dpf Tg[Gad1b:GFP; Vglut2a:DsRed] control (A_1-3_) (*N* = 8) and *scn1Lab* (B_1-3_) (*N* = 8) living larvae. (A_1_, B_1_) Inhibitory neurons (Gad1b:GFP) in 3 dpf control (A_1_) and *scn1Lab* (B_1_) larvae. (A_2_, B_2_) Excitatory neurons (Vglut2a:DsRed) in 3 dpf control (A_2_) and *scn1Lab* (B_2_) larvae. (A_3_, B_3_) Merged image of 3 dpf control (A_3_) and *scn1Lab* (B_3_) larvae. (C) Dorsal view schematic of 4 dpf larva. The red rectangle shows the region of interest. (E_1-3_, F_1-3_) Dorsal view of 4 dpf Tg[Gad1b:GFP; Vglut2a:DsRed] control (E_1-3_) (*N* = 8) and *scn1Lab* (F_1-3_) (*N* = 8) living larvae. (E_1_, F_1_) Inhibitory neurons (Gad1b:GFP) in 4 dpf control (E_1_) and *scn1Lab* (F_1_) larvae. (E_2_, F_2_) Excitatory neurons (Vglut2a:DsRed) in 4 dpf control (E_2_) and *scn1Lab* (F_2_) larvae. (E_3_, F_3_) Merged image of 4 dpf control (E_3_) and *scn1Lab* (F_3_) larvae. (H_1-3_, I_1-3_) Dorsal view of 5 dpf Tg[Gad1b:GFP; Vglut2a:DsRed] control (H_1-3_) (*N* = 8) and *scn1Lab* (I_1-3_) (*N* = 8) living larvae. (H_1_, I_1_) Inhibitory neurons (Gad1b:GFP) in 5 dpf control (H_1_) and *scn1Lab* (I_1_) larvae. (H_2_, I_2_) Excitatory neurons (Vglut2a:DsRed) in 5 dpf control (H_2_) and *scn1Lab* (I_2_) larvae. (H_3_, I_3_) Merged image of 5 dpf control (H_3_) and *scn1Lab* (I_3_) larvae. (D, G, J) Number of neurons in control and *scn1Lab* larvae at 3 dpf (D), 4 dpf (G) and 5 dpf (J). (K, L, M) Number of inhibitory and excitatory neurons in control and *scn1Lab* larvae at 3 dpf (K), 4 dpf (L) and 5 dpf (M). All images are representative 20 µm stacks acquired using LSM 880 Zeiss laser scanning confocal microscope equipped with a WPApo 20x/1.0 objective. Scale bar 40 µm. *N* = number of embryos. Error bars on all graphs represent standard error mean (SEM). *, *p* < 0.05; **, *p* < 0.01; ***, *p* < 0.001 indicate a statistically significant difference between control and *scn1Lab* larvae. Statistically significant difference between inhibitory and excitatory neuronal population is indicated as $$, *p* < 0.01; $$$, *p* < 0.001. *p*-values were determined using Student’s unpaired *t*-test.

### 3.4 Increased neuron apoptosis in Scn1Lab-depleted larvae

A large body of data gained from the analysis of neuronal tissues from human epileptic patients and animal epilepsy models, clearly indicates that both brief and prolonged seizures cause neuron death through activation of both apoptosis and autophagy signaling pathways [26]–[28]. We therefore investigated whether Scn1Lab depletion induced increased neuron apoptosis. We first used *in vivo* labeling with acridine orange. Results showed that the number of apoptotic cells was markedly increased in scn1Lab^AUG^ morphants compared to control larvae (Figure 4A, B, D; *p* < 0.001). To confirm this result, we also visualized neuron apoptosis using immunocytochemistry and anti-activated caspase-3 labeling. In good agreement with acridine orange staining, a significant increase in the number of activated caspase-3 positive cells were observed in the brain of Scn1Lab-depleted individuals compared to control larvae (Figure 4E, F, G; *p* < 0.001).

**Figure 4.**
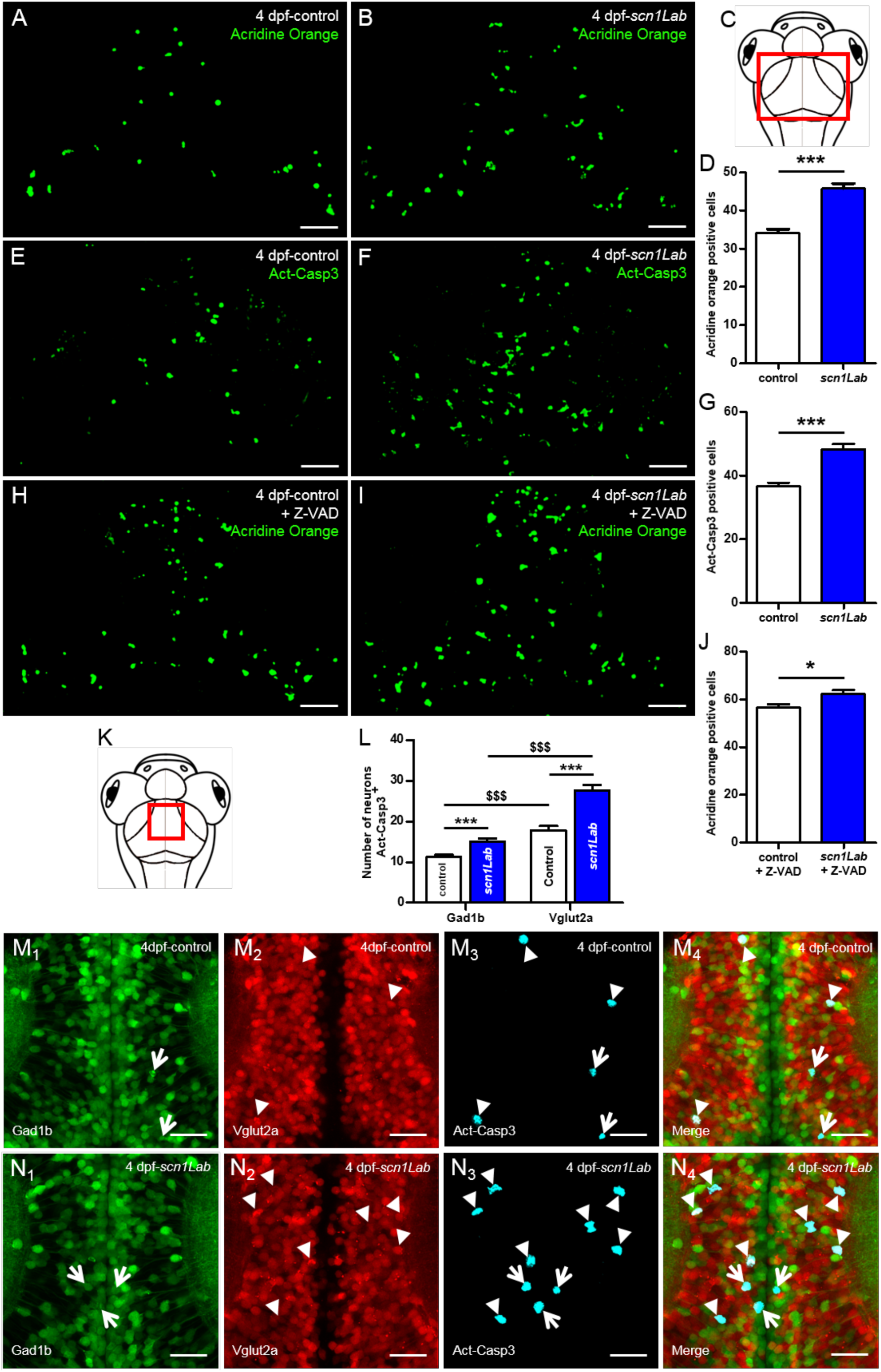
Increased neuronal death in Scn1Lab-depleted larvae. (A-B) Dorsal view of 4 dpf living control (A) (*N* = 15) and *scn1Lab* (B) (*N* = 15) larvae stained with acridine orange; 100 µm stack. Scale bar 40 µm. (C) Dorsal view schematic of 4 dpf larva. The red rectangle shows the region of interest for global cell death quantification. (D) Number of acridine orange-positive cells in 4 dpf brain. (E-F) Dorsal view of 4 dpf control (E) (*N* = 24) and *scn1Lab* (F) (*N* = 22) brain stained with activated caspase-3 antibody; 100 µm stack. Scale bar 40 µm. (G) Number of activated caspase-3-positive cells in 4 dpf brain. (H-I) Dorsal view of 4 dpf Z-VAD-treated control (H) (*N* = 18) and *scn1Lab* (I) (*N* = 16) larvae stained with acridine orange; 100 µm stack. Scale bar 40 µm. (J) Number of acridine orange-positive cells in 4 dpf brain. (K) Dorsal view schematic of 4 dpf larva. The red rectangle shows the region of interest for activated caspase-3 signal colocalization with inhibitory and excitatory neurons quantification. (L) Number of inhibitory (Gad1b) or excitatory (Vglut2a) neurons also showing activated caspase-3 labelling in control and *scn1Lab* larvae. (M_1-4_, N_1-4_) Dorsal view of 4 dpf Tg[Gad1b:GFP; Vglut2a:DsRed] control (M_1-4_) (*N* = 14) and *scn1Lab* (N_1-4_) (*N* = 14) brain stained with activated caspase-3 antibody; representative 15 µm stack. Scale bar 10 µm. (M_1_, N_1_) Inhibitory neurons (Gad1b:GFP) in control (M_1_) and *scn1Lab* (N_1_) larvae. (M_2_, N_2_) Excitatory neurons (Vglut2a:DsRed) in control (M_2_) and *scn1Lab* (N_2_) larvae. (M_3_, N_3_) Activated caspase-3 staining in control (M_3_) and *scn1Lab* (N_3_) larvae. (M_4_, N_4_) Merged image of control (M_4_) and *scn1Lab* (N_4_) larvae. Activated caspase-3 signal colocalization with excitatory neurons is shown by arrowheads where activated caspase-3 signal colocalization with inhibitory neurons is shown with simple arrows. Images were all acquired with an SP8 Leica laser scanning confocal microscope equipped with 40x/water 1.1 objective. *N* = number of embryos. Error bars on all graphs represent standard error mean (SEM). *, *p* < 0.05; ***, *p* < 0.001 indicate a statistically significant difference between control and scn1Lab larvae. A statistically significant difference between inhibitory and excitatory neuronal population is indicated as $$$, *p* < 0.001. *p*-values were determined using Student’s unpaired *t*-test.

Because Scn1Lab-depleted larvae showed an increased number of apoptotic cells, we next investigated whether this apoptosis was caspase-dependent. To test this hypothesis, we used Z-VAD, a cell-permeable caspase inhibitor that reversibly binds to the catalytic site of caspase proteases and inhibits their activity [29]. We incubated 3 dpf Scn1Lab-depleted and control embryos for 24 h in E3 medium containing 300 µM Z-VAD and 1% DMSO [30]–[32]. As negative control, age-matched Scn1Lab-depleted and control larvae were incubated for 24 h in E3 medium containing 1% DMSO. Cell death was assessed using acridine orange staining combined with real-time confocal imaging. Our data show that following Z-VAD treatment the number of acridine orange-positive cells was increased in both control (Figure 4A, H) and Scn1Lab-depleted larvae (Figure 4B, I), suggesting that Z-VAD had no effects on cell apoptosis in *scn1Lab* model.

We next examined whether apoptotic death occurred in excitatory or inhibitory neurons. To do this, 4 dpf Tg[Gad1b:GFP; Vglut2a:DsRed], Scn1Lab-depleted and Tg[Gad1b:GFP; Vglut2a:DsRed] control larvae, were fixed and analyzed by immunohistochemistry using anti-activated caspase-3 antibody. In both Scn1Lab-depleted larvae (Figure 4N_1_-_4_), and in control Tg[Gad1b:GFP; Vglut2a:DsRed] larvae (Figure 4M_1-_ 4), we observed a greater number of excitatory neurons expressing activated capsase-3 than inhibitory ones. Quantification of the number of inhibitory and excitatory neurons colocalizing with activated caspase-3 labelled cells, confirmed that neuronal death specifically affects excitatory neurons (Figure 4L; Gad1b^+^-Act-Casp3^+^ *vs* Vglut2a^+^-Act-Casp3^+^ controls: *p* < 0.001; Gad1b^+^-Act-Casp3^+^ *vs* Vglut2a^+^-Act-Casp3^+^ *scn1Lab*: *p* < 0.001).

## 4. Discussion

Dravet syndrome (DS) is a catastrophic childhood epilepsy with multiple comorbidities, which include severe intellectual disability, impaired social development, persistent drug-resistant seizures and high risk of sudden unexpected death in epilepsy (SUDEP) [1], [33]. In recent years, thanks to vertebrate genome conservation, full genome sequencing and readily available molecular tools and techniques, the zebrafish has become a powerful and versatile animal model for studying genetic forms of epilepsy and also to screen for novel anti-epileptic drugs [8], [10], [22]. Among epilepsy mutants, zebrafish larvae with loss of function of the *scn1Lab* gene, one of the two zebrafish orthologs of *SCN1A*, was instrumental to identify drugs alleviating seizures in refractory epilepsy [8], [10]. However, despite the large number of studies that have used Scn1Lab-depleted zebrafish mutants and morphants, the neuronal phenotypes induced by loss of function of this sodium channel encoding gene have not yet been fully investigated.

Using combined *in vivo* calcium imaging and local field potential (LFP) in the optic tectum of zebrafish models of DS, we observed a full correlation between calcium activity and neuronal activity, with both techniques consistently showing generalized ictal-like seizure events. Recent studies showed a correlation between calcium and local field potential activities using an acute PTZ seizure model in zebrafish [14]. To our knowledge, this is the first time that a correlation between calcium and electrical activities has been reported in a genetic model of epilepsy. This finding is important for research on epilepsy, because it validates the use of calcium activity to monitor the dynamics of seizures at the scale of an entire vertebrate brain. In particular, calcium imaging provides a unique opportunity to follow the dynamics of seizures spatiotemporally, from the initiation site to propagation through the brain.

We focused on excitatory and inhibitory synapses by studying two important proteins that are markers of synaptic integrity, PSD-95 and gephyrin [23]. PSD-95 by its PDZ domains serves as a scaffolding protein with a docking site for signaling proteins clustering around NMDA- and AMPA-type glutamate receptors, which are crucial for excitatory neurotransmission and synaptic plasticity [34]. Gephyrin is a core scaffolding protein, the postsynaptic component of inhibitory synapses, and it controls the formation and plasticity of inhibitory synapses through regulation of GABAA and glycine receptors [35]. In the present study, we demonstrated that the levels of PSD-95 and gephyrin are modified in zebrafish models of DS. Indeed, by studying the distribution of excitatory and inhibitory neurons during development, we have shown that at 3 days of development, the ratio of inhibitory to excitatory neurons was comparable between Scn1Lab-depleted larvae and their controls. Interestingly, from 4 days of development, we observed that the proportion of excitatory neurons increased when compared to that of inhibitory neurons in scn1Lab^AUG^ morphants, while this ratio remained unchanged in control embryos. This specific neuronal death may therefore explain the severe gephyrin signal loss observed in the *scn1Lab* model. Moreover, the increased in excitatory neurons detected from 4 dpf onward probably corresponds to the beginning of the disease in Scn1Lab-depleted embryos. Interestingly, these data are in good agreement with our results obtained par electrophysiological and calcium imaging, showing that the peak of seizures was observable at 4 and 5 dpf in Scn1Lab-depleted larvae [8].

Previous studies have shown that subtle changes in synaptic structure or composition, which alter synaptic plasticity and functional responses, most likely contribute to synaptic dysfunction [36]. Differences in key synaptic protein levels can provide information about the excitatory/inhibitory components of neurotransmission that might be disturbed or impaired during epiloptogenesis [37]. It is known that the balance between excitatory and inhibitory synapses is impaired in patients with epilepsy [20]. These results are also in line with the observation of Stief et al. (2007) in another model of epilepsy in rats, that the balance between excitatory and inhibitory synapses is shifted toward an over-excited state [19].

Our results using a zebrafish model of DS show an increase in apoptotic cells as previously observed in models of epilepsy [26]–[28]. We have also shown that treatment of Scn1Lab-depleted larvae with Z-VAD fails to inhibit cell death, suggesting that Z-VAD has no effects on cell apoptosis in *scn1Lab* model. However, we cannot exclude the possibility that Z-VAD may cause cellular toxicity in *scn1lab* model thus masking its protective effect. Unexpectedly, co-localization analysis of inhibitory and excitatory neurons with activated caspase-3 staining at 4 dpf revealed that cell death preferentially targeted excitatory neurons. This could be explained by the fact that during seizures, the excitatory neurons are hyper-excited and eventually die. In relation with this hypothesis, we sometimes observed during the *in vivo* calcium imaging using embryos from transgenic line Tg[HuC:GCaMP5G; *scn1Lab*] that some neurons became hyper-excited, as revealed by the over-expression of GFP labeling, and die (personal observations).

Our study emphasizes that the DS models are not only a powerful new tool for finding new antiepileptic drug candidates that could alleviate refractory seizures [8]–[10], but more importantly, they also display neurodevelopmental alterations characteristic of epileptic brains, such as the imbalance of excitatory /inhibitory synapses and apoptosis. Our data thus provide *in vivo* evidence for a developmental aspect of epileptogenesis that had been hypothesized [38], [39], but not definitely proved. Our study shows that DS is a disease of the central nervous system, and it calls for a closer focus on neurodevelopmental defects leading to severe comorbidities in DS (speech deficiency, autism, etc.) and not just on managing refractory seizures with antiepileptic drugs. More research is needed to understand the exact neurodevelopmental abnormalities causing epilepsy and to correct them as early as possible, rather than only treating them symptomatically at later stages. In the same way, we showed previously that gabra1 loss of function (generalized epilepsy) induced developmental alterations, in particular for the establishment of GABAergic networks throughout the brain [22].

The findings reported here shed light on the neurobiological changes that occur in the early stages of DS, and are a promising start in understanding the neurodevelopmental origins of DS and finding potential treatments to restore normal brain development at earlier stages of the disease.

## Supplementary Materials

Figure S1: Comparison of scn1Lab^AUG^ morphant and *scn1Lab^s552/s552^* mutant morphology and locomotor activity, Figure S2: Neuronal activity of *scn1Lab* model.

## Author Contributions

N.S.Y. and A.B., conceived and planned the experiments. A.B., performed the analysis, and designed the figures. R.H.A., and J.S., contributed to the analysis of the results. C.Y., was involved in the planning and the reviewing of the manuscript. N.S.Y., supervised the findings of this work, wrote and refined the article.

## Acknowledgments

We also thank Christiane Romain (Inserm UMR 1141), Olivier Bar (Inserm UMR 1141), and Solène Renault (Inserm UMR 1141) for their technical assistance.

