## Supplementary Figures for "Defective excitatory/inhibitory synaptic balance and increased neuron apoptosis in a zebrafish model of Dravet syndrome"

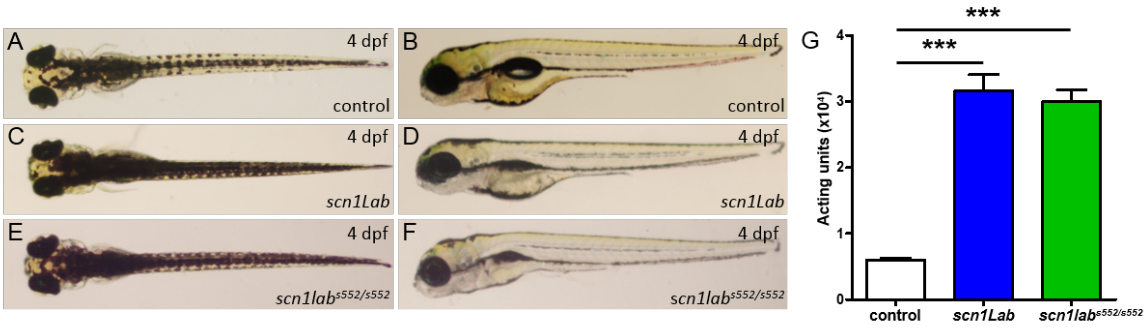


**Figure S1.** Comparison of *scn1Lab* morphant and *scn1Lab*$^{s552/s552}$ mutant morphology and locomotor activity. (A, C, E) Dorsal view of 4 dpf control (A), *scn1Lab* (C) and *scn1lab*$^{s552/s552}$ (E) larvae showing the hyperpigmentation of the morphant and the mutant larvae. (B, D, F) Lateral view of 4 dpf control (B), *scn1Lab* (D) and *scn1lab*$^{s552/s552}$ (F) larvae showing the impossibility for morphant and mutant larvae to inflate their swim bladder. (G) Plot of the locomotor activity of 4 dpf WT, *scn1Lab* and *scn1Lab*$^{s552/s552}$ larvae tracked during 25 min. Data were pooled from 5 independent experiments with at least 16 larvae per condition. Error bars on all graphs represent standard error mean (SEM). ***, $p < 0.001$. $p$-values were determined using Kruskal-Wallis test with Dunn's multiple comparison posttest.

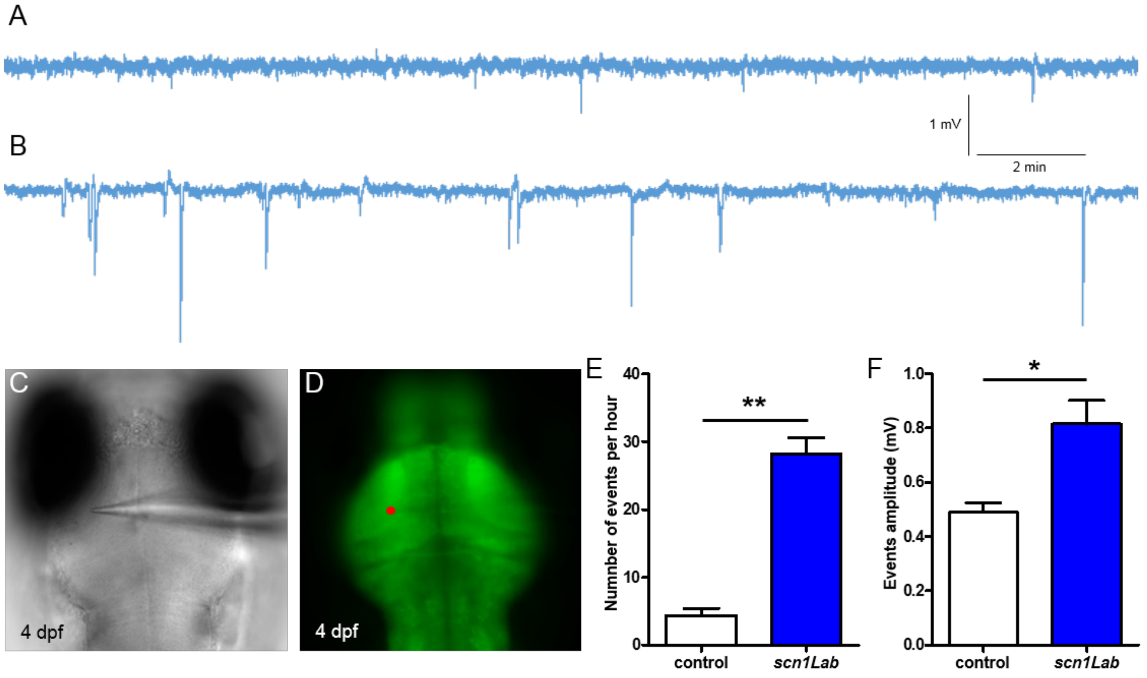


**Figure S2.** Neuronal activity of *scn1Lab* model. (A-B) Representative 20 min local field potential recordings in the neuropil of immobilized and paralyzed 4 dpf control (A) ($N = 5$) and *scn1Lab* (B) ($N = 5$) larvae. (C-D) Images of an immobilized and paralyzed larva during local field potential recording (C) showing the approximate localization of the electrode in the left neuropil using HuC-GCaMP5G transgenic line (D, red dot). (E) Number of events, downwards depolarization greater than 0.3 mV amplitude and lasting more than 100 s, during 1 hour recording. (F) Mean events amplitude. Error bars on all graphs represent standard error mean (SEM). *, $p < 0.05$; **, $p < 0.01$. $p$-values were determined using Mann Whitney test.
